# Tactile suppression during movement as optimal integration of somatosensory feedback across time

**DOI:** 10.64898/2026.02.25.707903

**Authors:** Fabian Tatai, Dimitris Voudouris, Dominik Straub, Katja Fiehler, Constantin A. Rothkopf

**Affiliations:** Centre for Cognitive Science, Technical University Darmstadt, 64283 Darmstadt, Germany; Institute of Psychology, Technical University Darmstadt, 64283 Darmstadt, Germany; Experimental Psychology, Justus Liebig University Giessen, 35394 Giessen, Germany; Computational and Biological Learning Lab, University of Cambridge, CB2 1PZ Cambridge, United Kingdom; Hessian Center for Artificial Intelligence, Darmstadt, Germany

**Keywords:** tactile suppression, optimal feedback control, attenuation, gating, feedback gains

## Abstract

Goal-directed movements generate tactile signals from the moving body, yet many are perceptually suppressed. Prevailing accounts propose that the nervous system reduces tactile sensitivity by prioritizing internal predictions over somatosensory feedback, contrasting with the view that suppression reflects peripheral masking. However, both accounts lack quantitative explanations of temporal dynamics and dependence on the motor task. Here, we show that tactile suppression is a consequence of optimal state estimation during movement. Using optimal feedback control theory, we derived how the nervous system should dynamically weight uncertain internal predictions against noisy somatosensory feedback. Human participants performed goal-directed reaching movements while vibrotactile stimuli were delivered at different time points. Suppression weakened as internal uncertainty about hand distance to target increased, consistent with greater reliance on sensory feedback. Model comparison identifies tactile suppression as dynamic, uncertainty-dependent state estimation rather than peripheral masking, establishing a normative principle for tactile sensitivity during action.

## Introduction

To move effectively, the brain must adequately process the continuously incoming sensory input to estimate the state of the body. Yet, when we move, the sensory signals from the moving body are suppressed, rendering external stimuli, such as the sleeve of a shirt sliding across the arm during reaching, less perceptible. Although this phenomenon has long been recognized, its functional role and underlying mechanisms remain debated [1, 2]. Importantly, tactile suppression during movement is not static but varies over time [3–10] and across motor tasks [5, 7, 11–16], suggesting that it reflects a flexible, motor-task-dependent process rather than a fixed reduction in sensitivity.

A central account, going back to Helmholtz in the 19th century [17], proposes that suppression arises from predictive mechanisms: expected sensory consequences of one’s own actions are suppressed via internal forward models [18] driven by efference copies of motor commands [19, 20]. Such models successfully explain why self-generated sensations, such as self tickling, are perceived as weaker [21]. However, they do not fully account for key features of tactile suppression during movement, including its temporal dynamics and its dependence on motor task demands. In parallel, alternative explanations attribute suppression to fixed gating mechanisms [1, 22]. One such account [23, 24] postulates that sensory signals generated by movement itself, i.e. the movement velocity, mask externally applied stimuli. While such mechanisms can contribute to suppression, they cannot readily explain why it varies with motor task [5, 7, 11–16] and why it arises prior to movement onset [9, 25]. Thus, a unifying account that explains both the temporal profile and motor task dependence of tactile suppression during movement is lacking.

Crucially, tactile signals are intertwined with motor action in a complex way [26]. Beside touch, they provide information that is directly relevant for estimating the state of the body as afferent signals from cutaneous mechanoreceptors encode kinesthetic information [27–30], like for example movement direction [31]. Disrupting cutaneous input impairs movement control [32], and artificial stimulation can bias perceived movement [33, 34], indicating that tactile afferents contribute to ongoing state estimation. These observations suggest that modulation of tactile processing during movement may reflect changes in how strongly such signals are weighted in the state estimation process.

Here, we propose that tactile suppression reflects body state estimation of hand distance to target under uncertainty. Specifically, we suggest that the nervous system dynamically modulates the processing of somatosensory feedback according to its relevance for estimating the current state of the body, similar as suggested for visual input during saccadic suppression [35]. This idea is grounded in optimal feedback control theory (OFC), which formalizes how noisy sensory feedback and uncertain motor predictions are integrated dependent on the motor task [36]. Within this framework, the relative contribution of sensory feedback is not fixed but varies over time, depending on the evolvement of the state estimate’s uncertainty due to signal-dependent motor noise. When internal predictions are reliable, sensory input can be down-weighted; conversely, when uncertainty of predictions increases, the system should rely more strongly on sensory feedback.

We tested this account by combining behavioral measurements of tactile detection during goal-directed reaching with computational modeling based on OFC with signal-dependent noise [37, 38]. Participants performed reaching movements while brief vibrotactile probes were applied to the moving limb at different time points. Detection performance served as a proxy for the processing of somatosensory feedback. Critically, we manipulated uncertainty about the initial state of the limb to test whether tactile suppression adapts accordingly.

Our results show that tactile suppression emerges well before movement onset and evolves dynamically throughout the movement, gradually recovering as the limb approaches the target. A computational model based on OFC explains the temporal profile of suppression during movement according to the change in uncertainty about predictions over future body state. As a crucial test of the model, we find that suppression is reduced when uncertainty about the initial state is increased, indicating a greater reliance on somatosensory feedback under these conditions. Model comparison demonstrates that alternative kinematic masking models can not explain the difference in suppression between uncertainty conditions. Furthermore, we report converging evidence of the model being in line with previously reported effects of task demands on tactile processing during reaching, when vision was removed.

Together, these findings suggest that tactile suppression is not a separate, non-specific gating mechanism, but rather reflects the dynamic regulation of sensory feedback in the service of state estimation. By linking tactile suppression to the principles of optimal feedback control, our work provides a unified account of its temporal dynamics and motor task dependence, and highlights the role of uncertainty in shaping sensory processing during action.

## Results

### Measuring tactile sensitivity during reaching movements

To investigate the time course of tactile feedback processing during movement, we carried out variants of the goal-directed reaching task, an experimental paradigm that has been extensively studied in the past [4–6, 13, 39]. We measured how well participants (*N* = 31) could detect near-threshold vibrotactile stimuli on their forearm during reaches to visual targets presented on a computer screen (Fig. 2a). Participants reached from a fixed starting position towards the screen, about 0.5*m* in front of them, at the center of which a circular target was displayed at various vertical positions. To dissociate suppression due to the predicted somatosensory input signals caused by touch at the moment of contact with the monitor (e.g., [7, 14]) and predicted somatosensory input arising because of the movement of a limb, we chose to probe tactile sensitivity at the forearm, where we expect less influence of the collision with the screen on tactile sensitivity [8]. Vibrations were administered with a frequency of 250*Hz* targeting the Pacinian corpuscles [40]. Besides their role in processing cutaneous touch, the Pacinian cells in the hand and lower leg have been shown to encode movement direction [27, 31], and it is therefore likely that they provide a similar function in the forearm. As they show a similar pattern of suppression to that of the muscle spindles [41], one of the main sources of kinesthetic sensory input, we reasoned that targeting cutaneous receptors with vibrations offers a behavioral method for measuring the time course of kinesthetic processing during movement. We adaptively delivered the stimuli to the reaching limb of individual participants, starting ∼ 250*ms* before movement initiation and continuing until shortly after the fingertip made contact with the screen. Participants completed 200 trials with a stimulus present, plus another 20 with no stimulus present, to measure false alarm rates.

**Figure 1.**
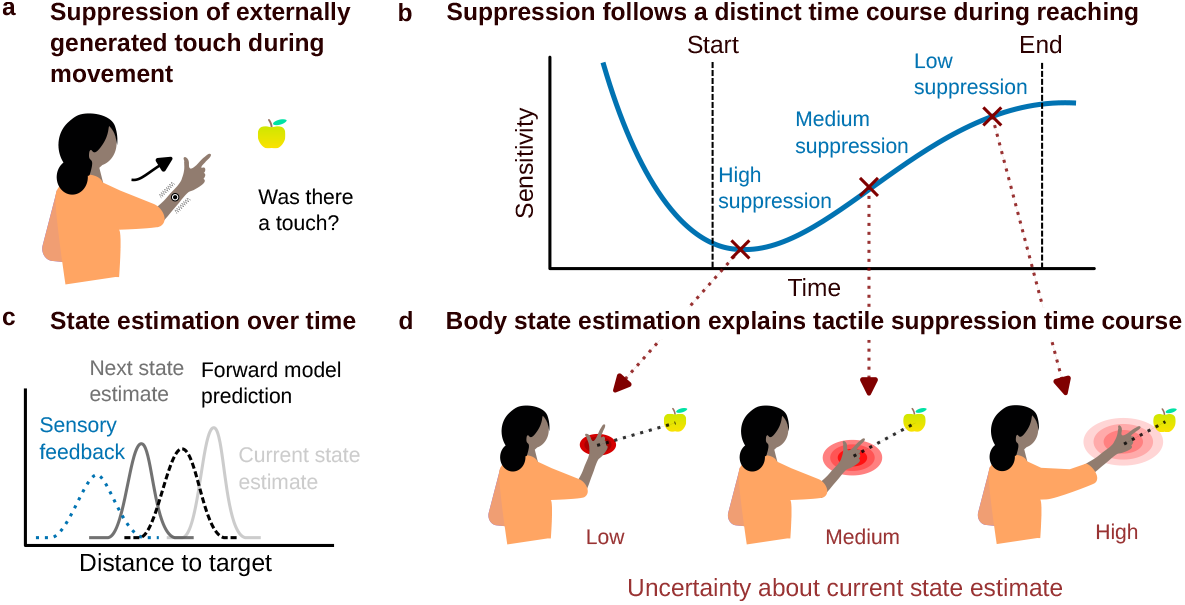
Suppression during movement: (a) tactile stimuli during movements, like goal-directed reaching, are suppressed on the moving limb. (b) This suppression is modulated over time, as demonstrated with an example time course reported in the literature [4, 5]. (c) During the movement, the nervous system needs to estimate the state of the body, for example, the hand’s distance to the target. It could do this by integrating the predictions of a forward model with sensory feedback over time. Note that during this dynamic process, uncertainties continuously interact and change the uncertainty about the current state estimate. Motor noise, for example, increases uncertainty about predictions. (d) When uncertainty about the state prediction is low, the system needs to rely less on sensory feedback and, thus, can suppress more sensory input. We propose that body state estimation of distance to target during movement explains the phenomenon of tactile suppression and its time course during goal-directed action.

**Figure 2.**
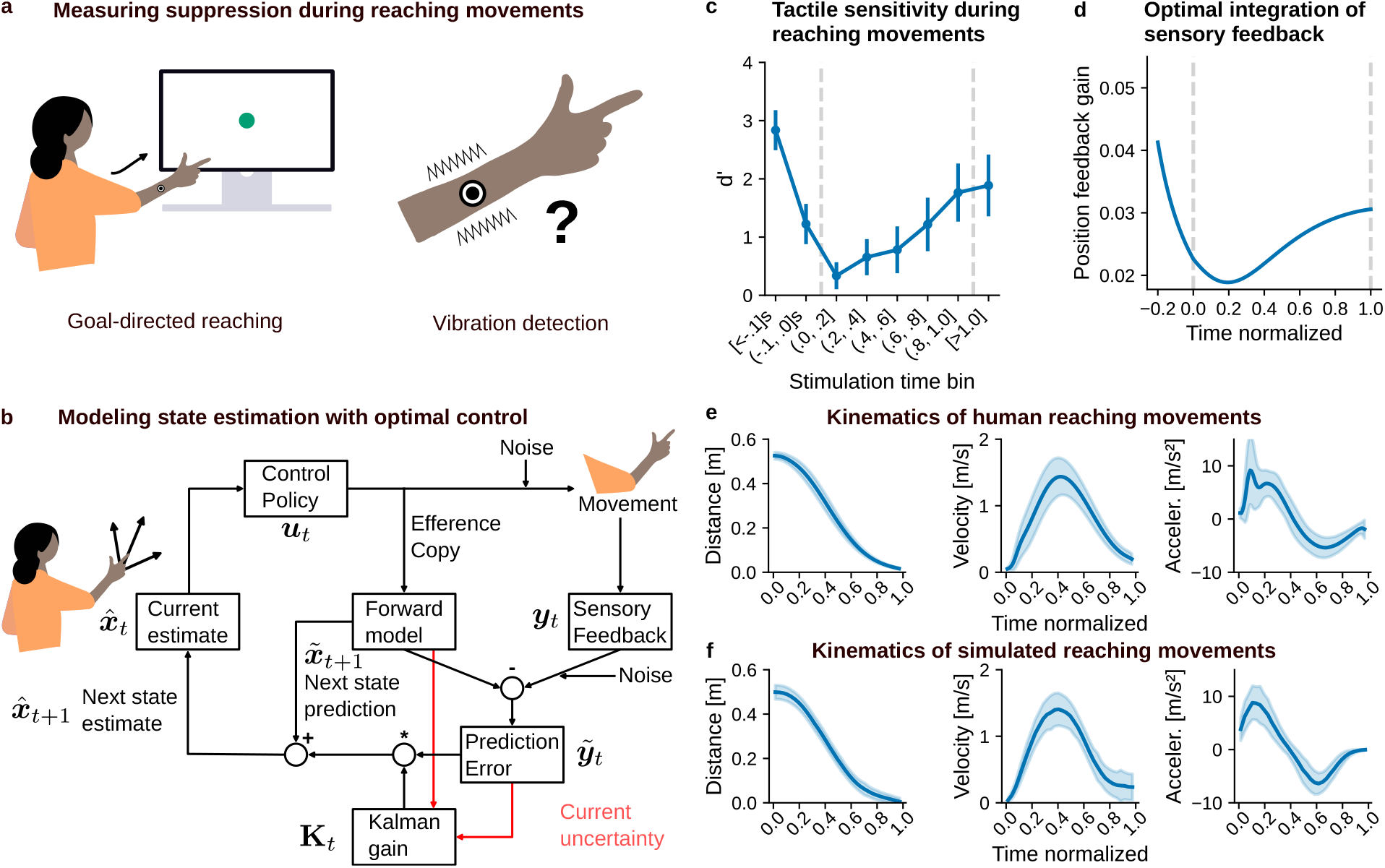
Measuring and modeling suppression of externally-generated touch during reaching movements. (a) Participants reached to a target on a computer screen, while they had to detect vibrotactile stimuli delivered at random in a time range from ∼ 0.25s before the movement onset until shortly after the finger reached the screen. (b) A model of reaching using optimal feedback control theory. Note that estimates of the system’s state are continuously updated over time. Dependent on the current certainty over the prediction and sensory feedback, the Kalman gain dynamically integrates sensory feedback with the forward model predictions. (c) Results: mean tactile sensitivity in *d*^′^ (with 95% confidence intervals) in time bins of 0.1s before movement start and in bins of 20% normalized movement time after movement start. Suppression starts well before the start of the movement (0s) and gradually recovers after 30% of the total movement time, though the recovery is not complete at the end. (d) The Kalman gain for sensory feedback about relative position over time resembles the pattern found in tactile sensitivity over time. (e) Mean and standard deviation of reaching kinematics across all participants and (f) the simulated kinematics of the model.

### Tactile sensitivity decreases initially and mostly recovers towards the end of the reaching movement

Fig. 2c shows the time course of tactile sensitivity during right-hand reaches, measured in *d*^′^, a standard measure in signal detection theory indicating the observer’s internal perception [42] (see Appendix A for criterion, and Appendix B for individual subjects). It can be interpreted as the distance between the noise and the signal plus noise distributions in units of their standard deviation. Higher values indicate higher sensitivity to the probe vibrotactile stimuli, whereas values close to 0 indicate almost no sensitivity. We binned stimulation trials according to the time point of their stimulation: [*<* −0.1]s and (−0.1, −0.0]s before movement onset, and after movement onset, normalized to the individual’s movement duration in bins of 20% of movement time until the movement end, and one bin for all stimuli delivered after movement end. The signal detection measures of *d*^′^ and the criterion *c* were computed per time bin and participant. We find that shortly before the start of the movement, sensitivity strongly decreases, and it begins to recover after ∼ 30% of the total movement time has elapsed. A repeated-measures ANOVA yielded a significant main effect of the stimulation time bin: *F*(2.894, 86.815) = 34.255, *p <* 0.001, 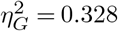 . Posthoc pairwise t-tests revealed that tactile sensitivity in the first 20% of the movement (0.0, 0.2] was worse than before movement initiation (*<* −0.1]s: *t*(30) = 13.345, *p <* 0.001, *d* = 3.125, *BF*_10_ *>* 100 (Bonferroni corrected). Furthermore, we find that at the end of the movement [*>* 1.0] sensitivity has not yet recovered to pre-movement level [*<* −0.1]s: *t*(30) = 3.973, *p* = 0.012, *d* = 0.779, *BF*_10_ = 72.27 (Bonferroni corrected).

### A model of sensorimotor integration

We use a model of single-joint reaching movements [38], which has been successful in explaining a wide range of empirical phenomena [43–46]. Specifically, reaching movements are modeled in the framework of OFC, which consists of a state estimator and a feedback controller [38]. Here, we focus on the state estimation part to explain the weighting of sensory signals relative to forward model predictions (see Optimal feedback control model of reaching movements for a complete description of the model).

The task-relevant state ***x***_*t*_ for the reaching consists of the distance of the hand to the target, its velocity and the force produced by the participant at time *t*. Participants receive noisy observations of this state *p*(***y***_*t*_|***x***_*t*_) = *N*(***x***_*t*_, **Σ**_***y***_). We assume that participants maintain a probabilistic estimate of the state, conditional on a sequence of sensory observations ***y***_1:*t*_ up to time *t*, in the form of a Gaussian probability distribution 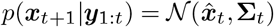, with a mean estimate 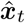 and a covariance matrix **Σ**_*t*_ representing the uncertainty about the state. At the beginning of the movement, this estimate is initialized with an initial covariance **Σ**_0_. As new sensory observations become available throughout the movement, the state estimate is updated by combining the predictions of an internal forward model with sensory feedback.

As we use a standard linear point-mass model of hand dynamics (see Appendix F), the mean prediction is given by linear state update equations

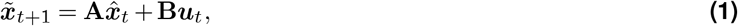

where ***u***_*t*_ is the participant’s control input, i.e. the muscle activations affecting the hand’s movement at time *t*. The matrices **A** and **B** describe the dynamics of the system, which models the translational movement of a point mass in a single dimension, a simplified model of decreasing the distance between the hand and the target by extending the elbow (see Appendix E and Appendix F for details). In our model we assume additive signal dependent normal motor noise and therefore the forward model predictions are also normally distributed 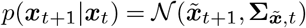 with prediction noise covariance 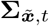 .

At each time step, participants receive a partial and noisy observation of the state ***y***_*t*_, which is integrated with the state prediction in an optimal fashion using the Kalman filter [37] to arrive at the next estimate of the state

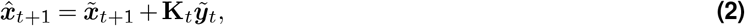

where 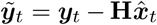 denotes the prediction error between the predicted sensory observation given the current state estimate and the actual observation (with linear observation matrix **H**). Consider that because of the noise in the prediction and the sensory noise the prediction error is also noisy 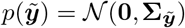 . The purpose of the Kalman **K**_**t**_ gain is to integrate the noisy sensory feedback signals about the state with the internal forward model’s noisy state predictions in a statistically optimal way by weighting the against each other: (Fig. 2b).

This weighting compares the uncertainties of the two signals 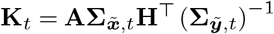 . As such, the Kalman gain can be related to the statistically optimal cue weights in cue integration [47, 48]. However, different from static cue integration paradigms, the two signals to be combined here are the somatosensory signals and the forward model’s prediction at each moment *t* in time. Because both signals’ uncertainties may change moment to moment, the relative weighting of the two signals by the Kalman gain may also change moment to moment. For example signal-dependent movement noise renders the forward model’s predictions more uncertain over time as the noise accumulates faster than sensory feedback can improve the accuracy of the state estimates which form the basis of the forward model predictions. Therefore, the dynamic evolution of the Kalman gain gives insight into how much somatosensory feedback should be weighted, i.e. suppressed, during the time course of a movement to achieve optimal inference of the state.

### Sensory feedback gains explain the time course of suppression

In order to relate the OFC model to the behavioral data, we analyze the Kalman gain **K**_*t*_, which integrates forward model predictions with somatosensory feedback by dynamically weighting the uncertainty in the current prediction of the body’s state against the uncertainty in somatosensory feedback. When the uncertainty in sensory feedback is larger relative to the uncertainty in the internal prediction, the Kalman gain downweights the contribution of the more unreliable sensory signal to achieve optimal integration. Specifically, we consider the entry of the Kalman gain matrix *K*_pos_ of the relative hand position, which constitutes the main state variable quantifying the motor task goal of the reaching task, i.e. the distance to the target (Fig 2d). We compare the time course of the Kalman gain with the sensitivity to somatosensory feedback measured with the vibrotactile stimuli targeting the cutaneous touch receptors, mainly Pacinian cells [40], which have been shown to be involved in limb state estimation [27, 31].

We find that the gain about the relative position feedback resembles the pattern found in our vibration detection experiment. Around movement onset, body state estimates and, therefore, also predictions of the forward model are more precise. Thus, the model prescribes that the sensorimotor system should rely less on somatosensory feedback about the relative position, and thereby show strong suppression. But as the movement unfolds, uncertainty in internal predictions increases with increasing movement speed due to signal-dependent motor noise, and thus somatosensory feedback becomes relatively more reliable. This suggests that tactile suppression during reaching aligns with the demands of state estimation in the motor task through the integration of somatosensory feedback and forward model predictions. It also highlights that the integration of somatosensory feedback during movement is a continuous process, as the uncertainty about the body state estimate continuously changes through the interaction of sensory and motor noise. While this provides a computational account for the observed time course of tactile suppression during reaching, this does not yet provide direct evidence for an adaptive uncertainty aware integration of sensory feedback with a forward model, which we consider in the following additional experimental condition.

### Tactile suppression is weaker when the state estimate about relative position is more uncertain

To examine whether the predictive system and uncertainty dependent weighting of somatosensory feedback is task specific, we explicitly manipulated the statistical properties of the motor task [1]. Therefore, we make use of OFC which prescribes how state estimation should change when properties of the motor task are varied. We devised a second condition in the reaching experiment based on model predictions, in which uncertainty in the initial state estimate Σ_0_ of the relative position between hand and target was increased (Fig. 3a). More uncertain state estimates imply less accurate predictions of the future body state and, in turn, require greater reliance on somatosensory feedback.

**Figure 3.**
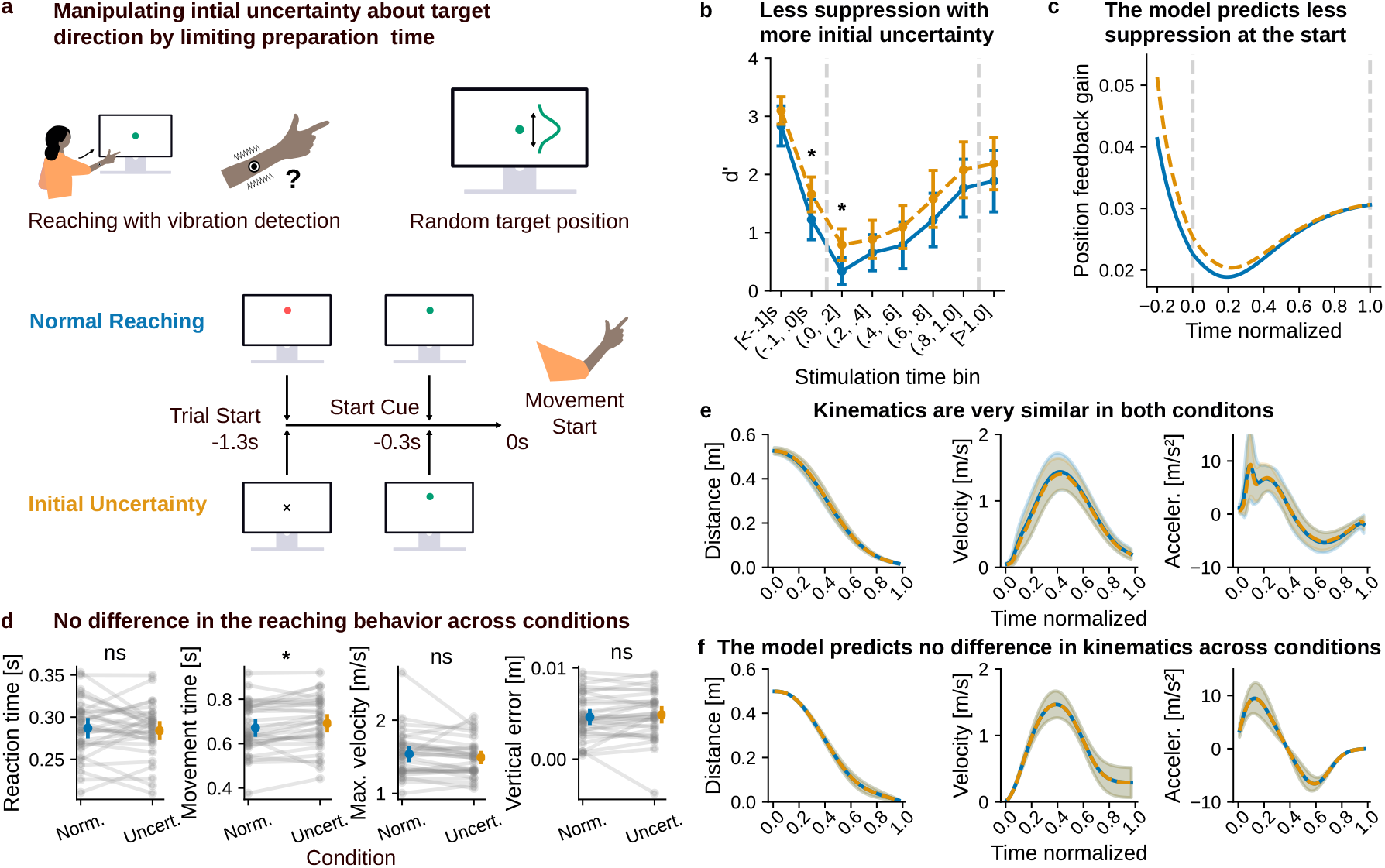
Manipulating initial uncertainty over relative position: (a) Participants detected vibrations during reaches to a target with varying vertical position on the screen. In the *certain* reaching condition, participants could view the target for a second, while in the *uncertain* condition, reaches had to be initiated right when the target was presented. By limiting the viewing time of the target, we increased participants’ initial uncertainty about their relative hand position to the target. (b) Participants suppressed less in the *uncertain* condition, especially around movement start. (c) This pattern between the conditions is predicted by the Kalman gain about the hand relative position in the optimal control model. (d) Mean descriptive statistics per participant; in color the per condition means and 95% confidence intervals. There was no or only very small significant difference between the conditions in any of the parameters, (e) as can also be seen in the mean and standard deviation of the kinematic timeseries and (f) as predicted by our model.

In the previously described *certain* condition, participants first saw the target, and a go-cue was given 1s later, requiring them to initiate their reach. In contrast, in the second *uncertain* condition, the target appeared at the same time as the go-cue. Therefore, in the *uncertain* condition, the same participants had to estimate their relative hand position while initiating the movement, whereas in the *certain* reaching condition, they had sufficient time to estimate the position before starting their reaching movement. Because of this, we assume that the participants’ initial uncertainty about the position of their hand relative to the target is increased in the *uncertain* condition. Therefore, based on the predictions of our model we expect weaker suppression in the *uncertain* condition, during the initial phase of the reaching movement (see Fig. 3c). Apart from this, the two conditions were identical.

We compared tactile suppression across the two conditions using a repeated measures ANOVA with the factors uncertainty condition and time bin. In line with the results reported above (not independent evidence as conditions overlap), tactile suppression was temporally modulated (main effect of time bin: *F*(2.550, 76.510) = 40.162, *p <* 0.001, 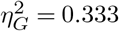). More importantly, and as predicted, tactile suppression was weaker in the *uncertain* than *certain* condition (*F*(1, 30) = 8.347, *p* = 0.007, 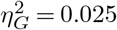 Fig. 3b). Although the interaction of uncertainty and stimulation time bin was not significant (*F* (4.889, 146.682) = 0.553, *p* = 0.732, 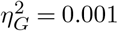), the model predicts that the difference between conditions should be largest around movement onset, where the initial uncertainty differs most, and should decline as the state estimate is refined over the course of the reach. Consistent with this, the reduction of suppression in the *uncertain* condition was strongest and most reliable around movement onset, surviving correction for multiple comparisons in the (−0.1, 0.0]s and (0.0, 0.2] bins (*t*(30) = 4.136, *p* = 0.002, *d* = 0.491, *BF*_10_ = 108.19; and *t*(30) = 3.535, *p* = 0.011, *d* = 0.656, *BF*_10_ = 25.30; both Bonferroni corrected), with decisive Bayesian evidence for a difference.

At the time bins after 0.2 the differences were numerically smaller and did not survive correction, and the corresponding Bayes factors were inconclusive (0.6 ≤ *BF*_10_ ≤ 1.4; for more details see Appendix C), indicating that our data can not determine whether the difference vanishes towards the end of the reach, as the model predicts, or persists at a reduced level. We therefore conclude that increasing the initial uncertainty reliably weakened tactile suppression around movement onset, in line with the predictions of the optimal control model, while a small residual difference across the reach cannot be excluded. Such a residual would be consistent with initial uncertainty that is not fully resolved during the beginning of the movement, for example because a fully specified movement plan, which can serve as a prior for the forward model, is not yet available in the *uncertain* condition, potentially pointing to sources of uncertainty beyond the single initial term Σ_0_ varied here.

The different time courses of sensitivity across the two conditions (*certain, uncertain*) might also be due to a difference in the underlying motor control strategy. However, a comparison of the kinematics makes this explanation rather unlikely (see Fig 3d). There were no significant differences between the conditions in reaction time (*t*(30) = 0.65, *p* = 0.523, *d* = 0.089, *BF*_10_ = 0.23), maximum velocity (*t*(30) = 1.38, *p* = 0.177, *d* = 0.166, *BF*_10_ = 0.45) and vertical endpoint error (*t*(30) = −0.98, *p* = 0.337, *d* = 0.096, *BF*_10_ = 0.30). There was a negligible but systematic difference in movement time (*t*(30) = −2.36, *p* = 0.025, *d* = 0.17, *BF*_10_ = 2.08), because the movement in the *certain* condition was just 20ms shorter on average than in the *uncertain* condition (see Appendix D for an overview over kinematic summary statistics). The mean positional, speed, and acceleration time series also appear similar between the *certain* and *uncertain* conditions (Fig. 3e).

These findings are also consistent with our model, which does not predict any effect of the initial uncertainty on movement kinematics (Fig. 3f). The intuition behind this is that the approximate relative position is available to participants at the start of the reaching movement, since the movement will always end up on the screen, and only its vertical component needs to be estimated more precisely. As the integration of sensory feedback is a continuous process, acquiring a more precise estimate of the relative position online is enough. To enable this, participants need to rely more on somatosensory feedback at the beginning of the movement because their current state estimates and predictions are more uncertain. Therefore, the increased uncertainty in the initial body state estimate can be resolved without changing the control of the reaching movement, while still providing the same precision at the end of the reach. Overall, the behavioral findings show that participants adjusted their tactile feedback processing based on the task demands without changing their kinematic control. This is in line with the model predictions and provides further evidence that tactile suppression is related to the estimation of the body state in the motor task.

### Optimal estimation, but not kinematic masking, explains the uncertainty dependence of suppression

A prominent alternative to a predictive account are fixed gating mechanisms. One such mechanism could be peripheral kinematic masking: the externally applied vibrotactile stimulus is rendered less detectable by the barrage of self-generated tactile and proprioceptive signals produced by the moving limb [9]. Masking is a bottom-up consequence of the movement itself, and it has been suggested that the resulting suppression should track movement kinematics, peaking near peak velocity [23, 24]. To our knowledge this account has only been demonstrated empirically but has not been modeled quantitatively. Therefore, we test a quantitative version, in which suppression is a function of the participant’s own reach kinematics:

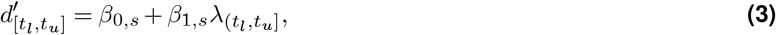

where *d*^′^ is the sensitivity, *λ* is the mean kinematic signal measured in the time interval (*t*_*l*_ : *t*_*u*_], and per subject offset *β*_0,*s*_ and slope *β*_1,*s*_. As candidate signals we consider velocity, but also acceleration and a linear combination of the two. But as the effect of kinematic signals might propagate across time, i.e. the current signal might mask stimuli before (backward masking) and after (forward masking) the current time point, we additionally considered filtered kinematic signals. Because forward and backward tactile masking differ in strength, with backward masking being stronger and longer lasting than forward masking [49], we modeled the masking signal as each participant’s velocity convolved with an asymmetric exponential kernel (see Kinematic masking models and model comparison). The values of this kernel were fixed to reflect the time course found by [49] (Fig. 4a). This timing also approximately aligns with movement related reafference reaching the brain [50, 51]. We additionally tested a symmetric kernel as a further baseline.

**Figure 4.**
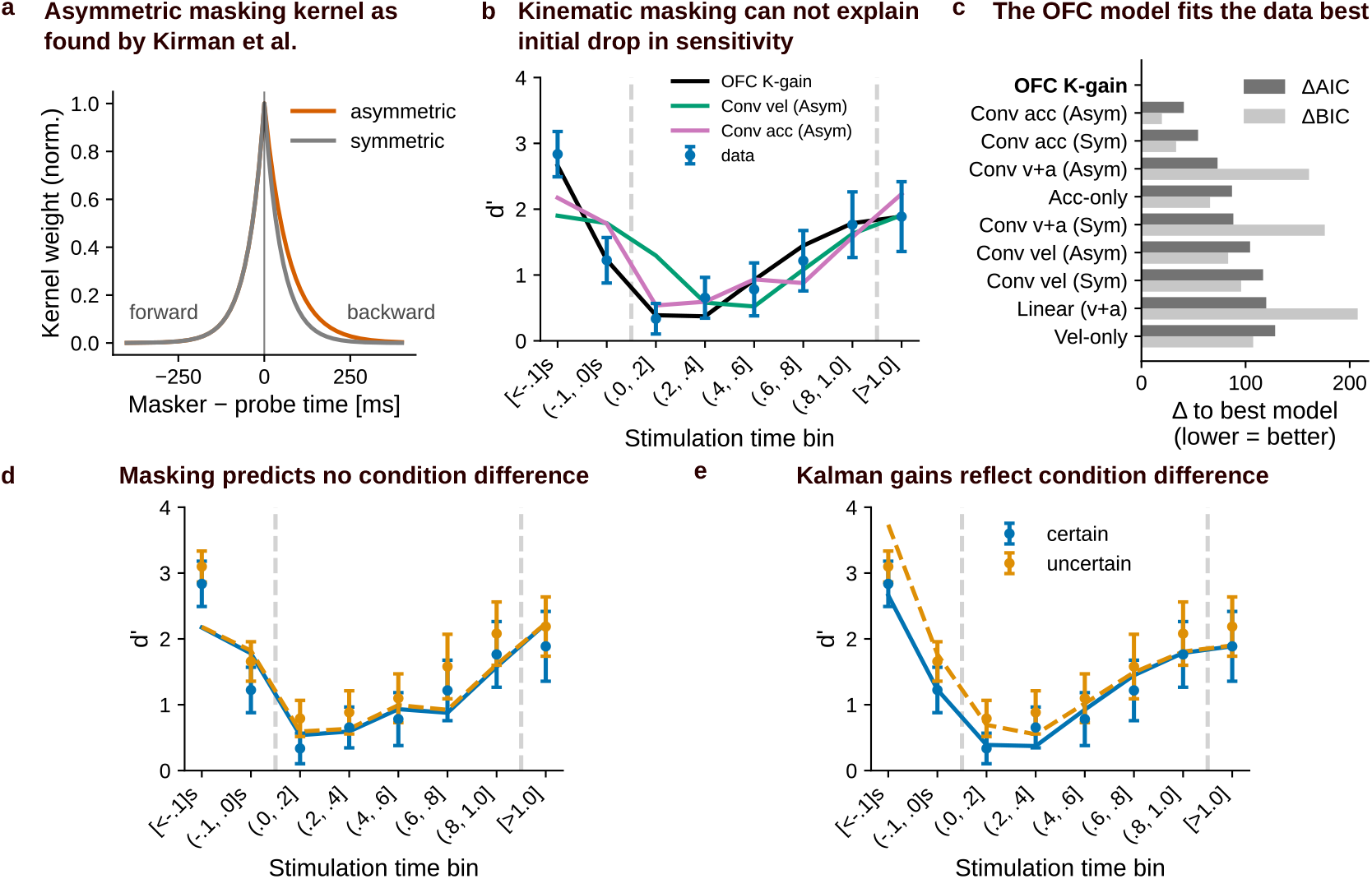
Optimal estimation, not kinematic masking, explains the pre-movement drop and the uncertainty effect. a) Fixed-shape masking kernels as a function of (masker − stimulus) time: an asymmetric exponential with a heavier backward (right) tail, following the relative strength of backward over forward masking reported by [49], and a symmetric exponential control. b)*certain d*^′^ time course (mean with 95% confidence interval) with the predictions of OFC model and the velocity and acceleration asymmetric-masking models. All models capture the post-onset recovery. Only the OFC model reproduces the drop before movement onset. c) Model comparison on the *uncertain* condition: ΔAIC and ΔBIC relative to the best model (lower is better). The OFC Kalman gain model fits best. d) Masking generalization: the fit of the link in the *certain* condition acceleration masking model applied to each condition’s own kinematics predicts no difference between *certain* and *uncertain*. e) Optimal-control generalization: the *certain* fit link applied to each condition’s Kalman gain predicts the observed reduction of suppression in the *uncertain* condition around movement onset.

In order to compare the masking model with optimal feedback control we consider Kalman gain as further candidate signal: 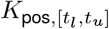 is the mean Kalman gain over relative hand position in the time interval [*t*_*l*_, *t*_*u*_]. Per participant Kalman gains were all computed with the same fixed parameters, only movement time was adjusted to the individual subjects movement time. All regression models thus have the same number of free parameters per participant (except for the combined acceleration and velocity models), so their fits are directly comparable by information criteria. Even though the parameters of the OFC model are not fit, we add additional 6 free parameters to the (AIC and BIC metrics) for the Kalman gain, as identified in Appendix H, for a conservative comparison.

During the movement, every model captures the gradual recovery of sensitivity about equally well, and the kinematic masking models reach a respectable fit (velocity- and acceleration-based models explain up to *R*^2^ = 0.735 of the baseline time course). The optimal control model nonetheless provides the best account overall explaining *R*^2^ = 0.759, with an AIC lower than the best kinematic model by 40.6 (Fig. 4c, Appendix I). Contrary to previous assumption [23, 24] models based solely based on velocity perform worst and acceleration seems more likely as a masking signal.

Critically, the masking models diverge at one point of the time course from the OFC model: the drop in sensitivity that occurs before movement onset (Fig. 4b). No kinematic masking model reproduces this drop, because there is essentially no movement, and hence no masking signal, before the reach begins. This is not an artifact of the chosen kernel: the asymmetric, backward-weighted kernel is precisely the variant most able to produce pre-movement suppression, since an imminent movement can mask a preceding stimulus, yet it still falls short because the velocity at onset is near zero. The optimal control model, by contrast, predicts the pre-movement drop directly from the decline of the Kalman gain in anticipation of the movement.

The central comparison, between the models, however, comes from the difference in the two conditions. The *uncertain* condition was designed a priori from the optimal control model’s prediction that increasing the initial state uncertainty Σ_0_ should weaken suppression around movement onset; the qualitative effect of increase in Σ_0_ was fixed from this design, only the amount of increase was set by hand. To test generalization, we took each model’s regression fit on the *certain* condition and applied it, unchanged, to the *uncertain* condition kinematics or Kalman gains. Because the reach kinematics are statistically matched across the two conditions (Fig. 3e), every kinematic masking model predicts essentially no difference between conditions (see Appendix I), for any kernel or parameter choice (difference before movement onset in time bin (−0.1,.0]s: Δ*d*^′^ ≤ 0.05; Fig. 4d). A model that depends only on kinematics cannot distinguish conditions whose kinematics are near identical. The optimal control model instead predicts a reduction in suppression at movement onset (Fig. 4e), reproducing the direction and time course of the observed effect from a link fit only to the baseline condition (predicted mean in time bin (−0.1, 0.0]s: Δ*d*^′^ = 0.54; versus an observed difference: Δ*d*^′^ = 0.43). Note, however, that the optimal control model overestimates the initial difference in bin [*<* .1]s (predicted mean in time bin (*<* −.1]s: Δ*d*^′^ = 1.07; versus an observed difference: Δ*d*^′^ = 0.26). This might be because the relationship of d’ and the Kalman gain contrary to our assumption is non-linear considering that d’ reaches a practical ceiling between 4 and 5.

Together, these analyses yield two independent dissociations that none of our kinematic masking models can produce, but that follow from optimal state estimation: the pre-movement drop, which masking cannot generate because the movement has not yet begun, and the uncertainty-dependent difference between conditions, which masking cannot generate because the kinematics are matched. Masking and optimal estimation make similar predictions during the movement, so peripheral masking may well contribute to suppression once the limb is in motion. Additionally, there could be masking model specifications explaining the pre-movement drop that we did not consider here. Masking because of pre-movement muscle activation [52], or a fixed centrally gated mechanism [1], could be potential explanations. We did not test these accounts, as to our knowledge no quantitative implementations of them exist, and in the case of pre-movement muscle activation we lacked the necessary measures. Crucially, though, none of these alternative mechanisms could explain the difference between the two conditions, which is our main dissociation.

### Tactile suppression is stronger when sensory feedback about target distance is more uncertain

The Kalman gain weights the forward-model prediction against the sensory feedback, and so depends on two distinct uncertainties: that of the prediction 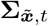 and that of the prediction error 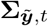, which in turn depends on the noise of the sensory feedback **Σ**_***y***_ (Fig. 2b). They enter the gain in opposite ways: raising the uncertainty of the initial state estimate and consequently the uncertainty of predictions, as in our *uncertain* condition, increases the gain and predicts weaker suppression, whereas raising the uncertainty of the sensory feedback lowers the gain and predicts stronger suppression. Essentially, one should trust sensory feedback less compared to forward model predictions when the feedback is less precise. Having tested the first prediction directly, we asked whether an independent data set is consistent with the second.

One way to reduce the quality of the sensory feedback, would be to remove visual information. The task-relevant state is the hand position relative to the visual target, and estimating it is inherently multisensory, requiring both the target and the hand to be localized in a common frame that vision normally provides [53, 54]. Only being able to access the goal position from memory, should lead to an overall increase in relative positional uncertainty [53]. As subjects could view the target before the movement, we assume this does not affect the initial estimate of relative position, but only subsequent sensory feedback arriving continuously during movement. Consequently, blind reaches to visually remembered locations cause significant errors in reach endpoint [54]. Additionally, previous research on the interaction of vision and tactile processing suggests that vision calibrates the tactile system, i.e., that the spatial and temporal precision of tactile feedback is better with vision [55–57]. Based on these collective findings, we assume that removing vision increases uncertainty of the sensory feedback about relative target position, which based on OFC should increase suppression.

Colino et al. [5] conducted a similar experiment to ours (Fig. 5a), where they measured tactile suppression during reaching to grasp movements. Although they measured suppression at multiple stimulation sites, in the following, we consider the right forearm, as it is most comparable to our experimental design. In their *certain* reaching-to-grasp condition (Fig. 5b), where the position of the grasping target is visible well in advance, they report a temporal modulation of tactile sensitivity that is similar to what we report in our current *certain* condition. Additionally, they included a *no-vision* condition, where participants’ vision was removed 500ms before the GO-cue. Here, tactile sensitivity was suppressed to a larger extent than when vision was available *F*(1, 9) = 6.03, *p* = .036 (as reported in the original work [5])(see Fig. 5b). In other words, participants showed stronger suppression when sensory uncertainty was higher. These findings in the *no-vision* condition align with the predictions of the OFC model (see Fig.5c) providing converging evidence for the idea that the dynamic weighting of sensory feedback and forward model predictions is adaptively adjusted to the demands of the motor task. Furthermore, they demonstrate how OFC can help interpret the effects of experimental manipulations reported in the literature.

**Figure 5.**
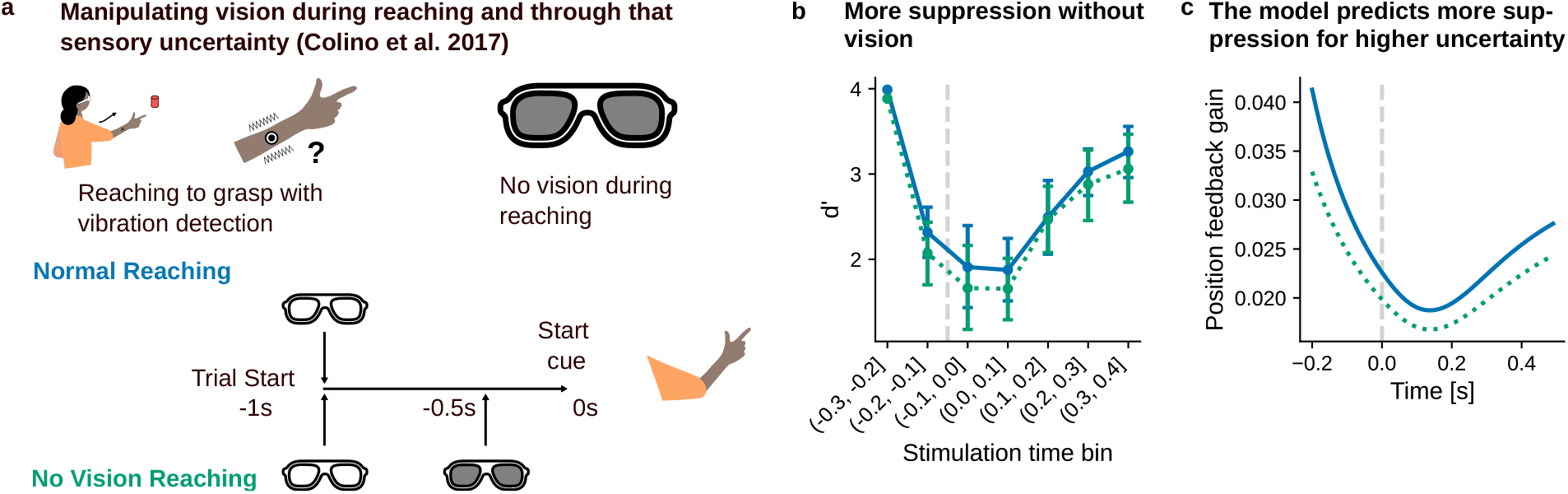
Manipulating uncertainty of sensory feedback about relative position by removing visual feedback: a) Colino et al. [5] conducted a similar study to ours, but ran a condition where vision was removed shortly before subjects initiated a reaching-to-grasp movement. b) They found that tactile sensitivity is reduced during the movement without vision. (digitized version of their plot) c) By understanding this manipulation as an increase in uncertainty of sensory feedback about relative position, our model accounts for the changed time course.

## Discussion

We show that tactile suppression during movement is a dynamic process shaped by the optimal integration of forward model predictions and somatosensory feedback, modulated by uncertainty about body state, and not non-specific gating mechanisms. Our results reveal that the time course of tactile suppression during reaching can be quantitatively explained by optimal feedback control theory that continuously integrates uncertain predictions with noisy sensory feedback. This normative framework accounts for our behavioral findings, and predicts how suppression adapts to changes in uncertainty over body state and sensory feedback, highlighting a general principle of dynamic processing of sensory information during movement.

Our study demonstrates that tactile sensitivity is temporally modulated during movement, with suppression weakening as uncertainty about relative limb position increases. Previous work has described this temporal modulation [3–10], but its underlying mechanism remained speculative. We propose that tactile suppression reflects the dynamic weighting of somatosensory feedback, such that suppression is reduced when the system relies more on sensory input, when uncertainty about body state is high. This idea is supported by our experimental manipulation of initial uncertainty, which increased reliance on positional feedback and altered the time course of suppression accordingly.

OFC provides a quantitative account of these effects, as it prescribes that the nervous system should weight predictions and feedback based on their respective uncertainties [37, 38]. Our model, which integrates forward model predictions with somatosensory feedback, successfully predicts the time course of suppression and its modulation by uncertainty and visual availability. This aligns with behavioral evidence for predictive mechanisms in tactile suppression [3, 11, 25, 58], and contrasts with accounts based solely on fixed gating [1, 2, 22].

Our computational account makes testable predictions for future research. Compared to other models, such as active inference [59, 60] or causal inference [61], OFC explicitly accounts for how uncertain sensory feedback and motor noise interact with the goals of the movement task. One such prediction is that the system should rely more on internal predictions when the quality of sensory feedback is diminished. In line with this, we report converging evidence based on past findings from Colino et al. [5]. They found increased suppression during reaching movements, when vision was removed, which degrades the quality of sensory feedback about hand distance to a visual target. Additionally, Colino et al. [4, 5] report an important control condition: they found no suppression on a stimulation site, not relevant for position estimation during a reaching movement (the other arm). A further possible application is that interactions between predicted sensations of expected touch and state estimation could explain the time course of suppression during reaching to one’s own hand [3, 62] or in reach-to-grasp movements [12].

While non-specific masking mechanisms may contribute at the implementational level [7, 22, 49], our model suggests that, from a computational perspective, the nervous system adaptively adjusts sensory feedback integration to the demands of the motor task. This interpretation is supported by our model comparison, in which we find that alternative kinematic masking models cannot account for the pre movement drop of suppression, and the difference in suppression between our uncertainty conditions. There are conceivable additions to the masking models, such as masking from pre-movement muscle activation [52] or a centrally triggered gate [1], which could possibly explain the pre-movement drop. However, the difference between the uncertainty conditions remains a crucial differentiator between OFC and the gating models.

Tactile suppression is often studied in the context of cutaneous touch, but our results highlight the role of body state estimation in the movement task. We assume that measuring sensitivity to vibrotactile stimuli during movement probes how much feedback from cutaneous mechanoreceptors is integrated for online control. This is supported by evidence implicating cutaneous touch receptors in state estimation [27–34]. As similar suppression patterns can also be observed in the muscle spindle afferents [41], our approach could inform neurophysiological experiments aiming to directly measure sensory feedback processing during movement.

Our findings have implications for understanding the neural basis of sensorimotor control. The cerebellum, which tracks body state and implements forward models [63–66], is a likely candidate for the uncertainty-dependent integration we describe. Neurophysiological evidence from monkeys shows top-down suppression of sensory input during movement [67], consistent with our model’s predictions. Moreover, our work suggests that bodily state representations include a notion of uncertainty, though how this uncertainty is represented in the brain remains debated. [68, 69].

Finally, by formalizing tactile suppression as an optimal integration process, we provide a quantitative framework for understanding how the nervous system regulates sensory input to optimize movement. This perspective not only advances our understanding of sensorimotor control, but also opens avenues for clinical applications—e.g., in prosthetics [70] and neuropsychiatric disorders where predictive mechanisms may be impaired [65, 71].

## Methods

### Participants

31 undergraduate students at the Justus Liebig University Gießen joined the study (19 − 34 years old, mean age: 26; 21 females, 10 males). They were all naive with regard to the goals of the study. They received either course credits or an hourly compensation of 8 EUR for their effort. Participants were right-handed according to the German translation of the Edinburgh Handedness Inventory (EHI: 88, range: 37 − 100; [72]) and had normal or corrected vision. The experiment was approved by the Ethics Committee of Justus Liebig University Giessen, and all participants gave their written informed consent before joining the experiment.

### Experimental Setup and Apparatus

The experiment was conducted in a well-lit room. Participants sat in front of a table (118 × 80cm), facing a monitor (VPixx Technologies Inc., Saint Bruno, Canada; 1920 × 1080px; 521 × 230mm; 60Hz) placed at a distance of 52cm in front of them. We used this monitor to present visual stimuli that were important for the experiment. To probe tactile sensitivity, we delivered vibrotactile stimuli using a small actuator (diameter of 0.8cm; Engineer Acoustics Inc., Florida, US) housed within a plastic case (2.2 ×4.5 ×0.5cm), hereafter referred to as *tactor*. The tactor was fixed with medical tape to the dorsal part of the participants’ forearm, halfway between the elbow and wrist. We ensured that the tactor was fixed tightly enough so that it could not move during the experiment, but not so tightly that it would hinder blood flow and, thus, influence tactile sensitivity. In addition, the position of an infrared marker, fixed to the participants’ right index fingernail with adhesive putty, was recorded at 100 Hz with a single-camera Optotrak Certus system (Northern Digital, Waterloo, Canada). We fixed the cables of the tactor and the infrared marker along the participants’ right arm with tape to ensure that they would not hinder their arm movements, and to prevent any cable movement during the reaching task, as such movement could generate unintended tactile sensations.

### Procedure

The experimental procedure always started with a baseline tactile detection task performed at rest. After that, participants performed a practice reaching block, followed by two experimental blocks (one per reaching condition *certain* and *uncertain*) where the tactile task was performed during reaching movements. These two reaching blocks were presented in counterbalanced order across participants.

#### Baseline tactile detection task

The baseline task allowed us to first estimate each participant’s tactile detection threshold without any influences of movement planning or execution. This is important because tactile sensitivity is highly variable between individuals [73]. We used this baseline sensitivity to present tactile probes during subsequent reaching tasks tailored to each participant’s detection thresholds. Participants performed the baseline block with both hands resting on the table at a comfortable position. Each trial began with the participant pressing a keyboard button placed below their left index finger. A brief tactile stimulus (50ms, 250Hz) was delivered to the participant’s right forearm at a random time point drawn from a uniform distribution between 100 and 400ms after the button press. This tactile stimulus could have one of 9 possible intensities, with a peak-to-peak displacement ranging from 38 to 341*µ*m, in steps of 38*µ*m. Each of these intensities was presented 4 times. In addition, we included 20 trials without stimulation (catch-trials). This resulted in a total of 56 trials for the baseline task, presented in a random order. At the end of each trial, a question on the monitor prompted participants to respond whether they felt a tactile stimulus or not. The answers “Yes” and “No” appeared at two lateral positions on the monitor (counterbalanced across participants), and participants had to press one of two pedals with their feet to select the desired answer before they could start the next trial. Participants gave their response to the tactile task and initiated each trial at their own pace. The baseline block lasted ∼ 5 minutes.

Based on responses to the tactile stimuli, we estimated a tactile stimulus intensity, which served as the basis for the probe tactile stimulus in the subsequent reaching trials. Details about this are provided in Calculation of individual stimulus intensity.

#### Practice and Reaching trials

Immediately after the baseline task, participants performed 40 trials to practice their reaching movements. These were important for participants to get familiar with the task and for us to determine the meaningful period to probe tactile sensitivity (see below). Participants started each reaching trial with both hands resting palm-down on the table, with their right index finger above a button at the starting position. This position was 10cm in front of the participants, aligned with their midline and the center of the monitor. To begin a trial, participants had to press and hold this starting button, which immediately triggered the presentation of a red circle (diameter of 40px) at the horizontal midline of the monitor, while its vertical position was drawn from a uniform distribution spanning 300px above to 300px below the screen center. After a random delay between 700 and 1300ms, the red circle turned green, serving as a GO-cue for participants to reach and hit it. No tactile stimulus was delivered during these 40 trials. The practice block lasted ∼ 5 minutes.

Our main interest was whether tactile suppression depends on the uncertainty about the initial position of the hand relative to the target. To this end, the two experimental conditions differed in whether the target position was known to the participant before the GO-cue. In the *certain* condition, the target presentation followed the same procedure as the one described above for the *practice* block. Thus, participants knew the required movement direction well before the GO-cue was presented. In contrast, in the *uncertain* condition, pressing the start button did not reveal the target position. Instead, a black cross (10 × 10px) appeared centrally on the monitor. After a random interval between 700 and 1300ms, a green circle was shown at the target position. This target was identical in size and was drawn from the same vertical position distribution as in the *certain* condition. Therefore, both conditions involved the same possible range of target positions and differed only in the time point when the target position became known to the participants.

Participants had to start their reaching movement within 500ms after the presentation of the green target (GO-cue). If they released the button before or after this temporal window, a message appeared on the monitor warning them that they started too early or too late, respectively, and the trial was immediately repeated. We probed tactile sensitivity in the reaching trials by delivering a brief vibrotactile stimulus (50ms, 250Hz) of fixed intensity to the participants’ right forearm. The stimulus could be delivered at a random moment during the trial, within a specific temporal window. Specifically, for the first ten trials of both the *certain* and *uncertain* conditions, the probe stimulus was delivered at a random time drawn from a uniform distribution between 100 and 400ms after the presentation of the red circle or black cross, respectively. From the 11th trial onward, the timing of the probe was adjusted based on each participant’s recent reaction and movement times [12, 25, 73]. For each trial, we calculated the reaction time as the interval between the GO-cue and the release of the start button, and the movement time as the interval between movement onset and movement end. Movement time was determined based on online analysis of the kinematic data: movement onset was the first frame when the three-dimensional speed of the fingernail marker exceeded 5 cm/sec, and movement end was the first frame after movement onset when the marker speed dropped back below that threshold. From the 11th trial onward, we estimated the median reaction time and median movement time of the preceding 10 trials. The sum of these two medians provided an estimate of the expected duration of the upcoming trial. We delivered the probe tactile stimulus at a random time drawn from a uniform distribution between 100 ms after the GO-cue and this estimated duration. Each reaching block included 200 trials with a probe stimulus and another 20 trials without stimulation (catch-trials).

## Data Analysis

### Calculation of individual stimulus intensity

The responses to the tactile stimuli during the baseline task were fitted with a logistic function using maximum-likelihood estimation via the function `psignifit 3` [74] in MATLAB 2019b. From the resulting psychometric function, we determined the baseline tactile intensity as the stimulus intensity at 80% correct performance. For the reaching trials, we set the probe to a fixed stimulus intensity, allowing us to calculate sensitivity and criterion using signal detection analysis. The probe stimulus intensity in the reaching trials was set to three times the rounded-up baseline intensity. We chose to use as baseline sensitivity the intensity at 80% and to triple this value to achieve a relatively strong stimulus tailored to each participant’s sensitivity, thereby alleviating the strong suppression that is often observed on a moving limb [13, 73]. For this reason, whenever the tripled baseline intensity was smaller than 47*µ*m, the probe stimulus intensity was set to 47*µ*m.

### Kinematic analysis

From the collected motion capture data, we first calculated 3D velocity and acceleration by differentiating the positional data. The start of the reach was defined as the time point of lift-off, determined by the time of the start button release. The end of the reach was defined as the point in time when the velocity, after having passed the critical velocity of 0.1m*/*s once after lift-off, dropped below this critical value again. Subsequently, we excluded trials where we could detect no reach or when the endpoint error larger than 0.05m, which amounted to 51 trials. Based on the extracted reaches, we calculated the kinematic summary statistics (see Appendix D) and normalized the movement time. For the mean kinematic trajectories Figure 2e and Figure 3e we normalized in movement time before averaging, rather than binning across participants. Within each participant, every reach’s distance, velocity, and acceleration were interpolated onto the common normalized-movement-time grid and averaged across trials. We then plot the group mean and standard deviation of these per-participant profiles across participants.

### Signal detection analysis

As we delivered the vibrotactile stimuli adapted to the individual movement times, we decided to bin the data by stimulation time point normalized to the overall movement time, which varied within participants during the time course of the experiment, but also across participants. Stimuli delivered before movement onset were binned in absolute times. With this procedure, the time bins contained 25 trials on average per subject. Using the false alarm rate measured per condition and the hit rate per time bin, we calculated *d*^′^ and *c* with standard correction of extreme rates [75].

### Statistical testing

The obtained kinematic summary statistics as well as the signal detection parameters were submitted to statistical testing with the python package `pingouin` [76]. Mauchly’s test indicated violations of sphericity for the stimulation time bin in both ANOVAs (*W* = 0.011 and *W* = 0.004, both *p <* .001), though not for the interaction (*W* = 0.246, *p* = .072). Greenhouse–Geisser-corrected degrees of freedom and p-values are reported throughout; the two-level condition effect needs no correction. Cohen’s d for paired comparisons was computed with the pooled standard deviation of the two conditions.

### Optimal feedback control model of reaching movements

We modeled reaching movements using a simplified model of a single-joint arm movement in the framework of stochastic optimal control [38] (see Appendix E). The hand is modeled as a point mass. Its state 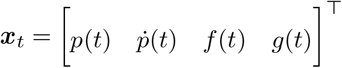 includes the hand’s relative position to the target (here modeled in 1D as the distance to target) *p*(*t*), velocity 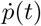, and force *f* (*t*), and an auxiliary state *g*(*t*) used to implement a muscle-like low-pass filter (see Appendix F for details). The temporal evolution of the state can be described with a linear dynamical system with signal-dependent noise

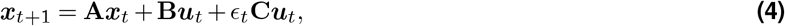

where *ϵ*_*t*_ ∼ *N* (0, 1). Due to the final term, the noise in the control input scales with its magnitude [77], such that faster movements requiring more force will show more variability. At each time step, the agent receives a noisy sensory observation of the state of the system

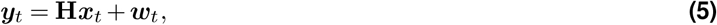

with ***w***_*t*_ ∼ *N*(**0, Σ**_***w***_). The observation matrix **H** is chosen such that the agent receives an observation of the hand’s current distance to the target, its velocity, and force (see Appendix F). Based on the sensory observations, the agent infers a posterior distribution about the state 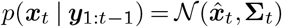 . The agent has initial uncertainty about the state, represented by 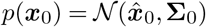, with

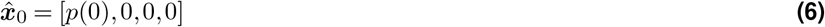

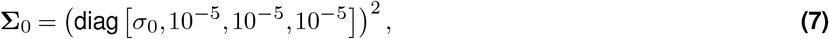

where *p*(0) is the intial distance to the target and *σ*_*o*_ the uncertainty about the distance. This distribution is updated using sequential Bayesian inference, which results in the Kalman filter [37] equations for the mean state estimate

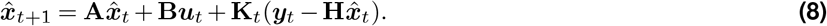

To derive **K**_*t*_, we use the generalized Kalman filter equations with signal-dependent noise [38]. For details and derivations, see Appendix E. We can decompose the state estimation problem into two steps. First, the agent uses an internal forward model of the dynamics to compute a prediction of the state 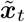 from their current state estimate 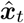 and the control input ***u***_*t*_:

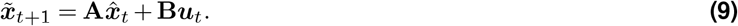

Second, the predicted state is updated using the difference between the noisy sensory observation and the predicted sensory observation 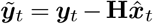, which let us write Eq. (8) in the following form

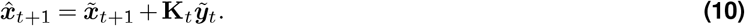

Intuitively, the Kalman gain **K**_*t*_ can be interpreted as a weight analogous to a cue weight in cue integration paradigms that determines the extent to which the current noisy sensory measurement is combined with the noisy prediction from the internal forward model [47, 48]. To achieve the statistically optimal estimate of the current state, a cue contributes more to the integrated estimate the more reliable it is, i.e., the lower its uncertainty.

The cost function

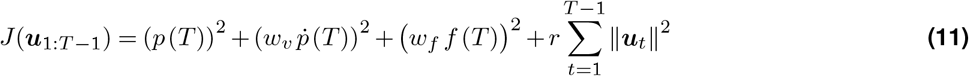

expresses the task of reaching the target (i.e. minimizing squared distance to the target) and stopping there (i.e. minimizing final velocity and force) at the final time step *T*, while minimizing the cost of control effort exerted along the way.

The optimal action that minimizes the expected cost can be computed as a linear function of the current state estimate

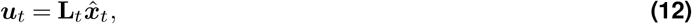

where the matrices **L**_*t*_ are called the control gains (see Appendix E). Using this process of optimal state estimation and control, we simulate the system dynamics (Eq. (4)) to compute a sequence of states ***x***_1:*T*_ . We can also inspect the sequence of Kalman gains **K**_1:*T*_ to investigate how sensory information was integrated over the course of the movement. Our implementation of the OFC model is based on a code framework published alongside previous work by some of the authors [78]. A detailed description of our model simulations can be found in Appendix G and see Appendix H for how the free parameters of our model influence the time course of the Kalman gain and how robust our reported effects are to variations of those parameters.

### Kinematic masking models and model comparison

To test whether tactile suppression is a fixed, bottom-up function of the participant’s own reach kinematics rather than a consequence of predictive state estimation, we fit a family of kinematic masking models to the certain-condition sensitivity time course and compared them against the optimal feedback control account on identical footing.

### Continuous kinematic profiles

For each participant we first constructed continuous velocity and unsigned acceleration profiles in real time (see Kinematic analysis). Each normalized profile was then stretched to that participant’s mean movement time, padded with rest (zeros) for 0.2 s before and after the reach, and resampled onto a shared real-time grid with step 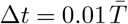, where 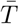 is the mean reach duration. All subsequent kinematic features, whether raw or filtered, were derived from these continuous profiles.

### Masking kernels

Because the effect of a kinematic signal can propagate across time (the signal at one instant may mask a stimulus presented before or after it), we additionally considered temporally filtered kinematics. Forward and backward tactile masking differ in strength, with backward masking (a masker following the stimulus) being stronger and longer lasting than forward masking [49]. We therefore convolved each kinematic profile with a fixed asymmetric exponential kernel as a function of masker–stimulus time *τ*,

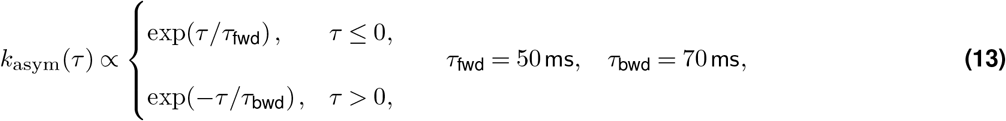

whose heavier backward (right) tail reflects the relative time course of backward over forward masking [49]. As a control we also used a symmetric exponential kernel,

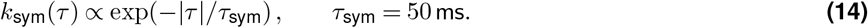

Both kernels were normalized to unit area before convolution, so that filtering redistributes but does not rescale the kinematic signal. The kernel shapes were fixed a priori to match the results from [49]. In addition, to peripheral backward masking the filter also approximately corresponds to the time it takes for movement related reafference to reach the brain [50, 51].

### Candidate signals

As candidate masking signals we considered the raw velocity, the raw unsigned acceleration, and their linear combination, as well as the velocity and unsigned acceleration each convolved with the asymmetric and with the symmetric kernel. As the optimal-control comparison we used the Kalman gain over relative hand position, *K*_pos_, obtained from the reaching model with fixed parameters and only the movement time adjusted to each participant.

### Binning and the sensitivity link

Each candidate signal was reduced to one value *λ* per stimulation time bin by averaging the (optionally filtered) continuous signal over the corresponding bin window, using the same binning as the detection data: absolute-time bins before movement onset, fractions of the participant’s movement time during the reach, and one post-movement bin. Every model was linked to sensitivity through the same per-participant linear regression, with an individual offset and slope,

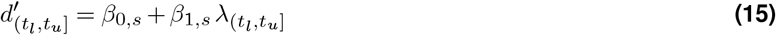

for the masking models, and the analogous link with *K*_pos_ for the optimal-control model. The regression was fit by ordinary least squares to the *certain* condition only. Because each model contributes one offset and one slope per participant, all models carry the same number of free parameters per participant, except the combined velocity-and-acceleration models, which carry one additional slope.

### Model comparison

We assessed the fits with the coefficient of determination *R*^2^, the mean squared error, and the Akaike and Bayesian information criteria (AIC and BIC), computed in the standard way for the per-participant linear regression. Because the Kalman gain is itself the output of the optimal control model, we conservatively added 6 free parameters to the optimal-control model when computing its AIC and BIC, so that the comparison does not favor it by ignoring the parameters of the reaching model (see Appendix H).

### Generalization across conditions

To test whether each model can reproduce the observed difference in suppression between conditions, we took the per-participant coefficients fit on the *certain* condition and applied them, unchanged, to each condition’s own signal: the participant’s own kinematics for the masking models, and the participant’s own Kalman gain for the optimal-control model. The predicted between-condition difference Δ*d*^′^ then follows from the signals alone, with no additional fitting.

## ACKNOWLEDGEMENTS

This work was supported by the Deutsche Forschungsgemeinschaft (German Research Foundation, DFG) under Germany’s Excellence Strategy EXC 3066/1 “The Adaptive Mind”, Project No. 533717223, by the European Research Council (ERC Consolidator Award “ACTOR”-project number ERC-CoG-101045783), and by the Collaborative Research Center SFB/TRR135, project A4, under grant agreement 222641018. We thank Malaika Grace Alphonsus for the support with data collection.

## AUTHOR CONTRIBUTIONS

Conceptualization, Methodology: all authors; Funding Acquisition, Resources: D.V., K.F., C.R; Investigation: D.V.; Data Curation: F.T., D.V.; Formal Analysis: F.T., D.S., C.R.; Software: F.T., D.V., D.S.; Supervision: K.F., C.R.; Validation: F.T., D.S.; Project Administration, Visualization, Writing - original draft: F.T.; Writing - review & editing: D.V.,D.S,K.F.,C.R.

## ETHICS STATEMENT

The experiment was approved by the Ethics Committee of Justus Liebig University Giessen, and all participants gave their written informed consent before joining the experiment.

## DATA AVAILABILITY STATEMENT

The data and code for this project is available here.

## Supplementary Information

### A Mean signal detection parameters across time

**Figure S1.**
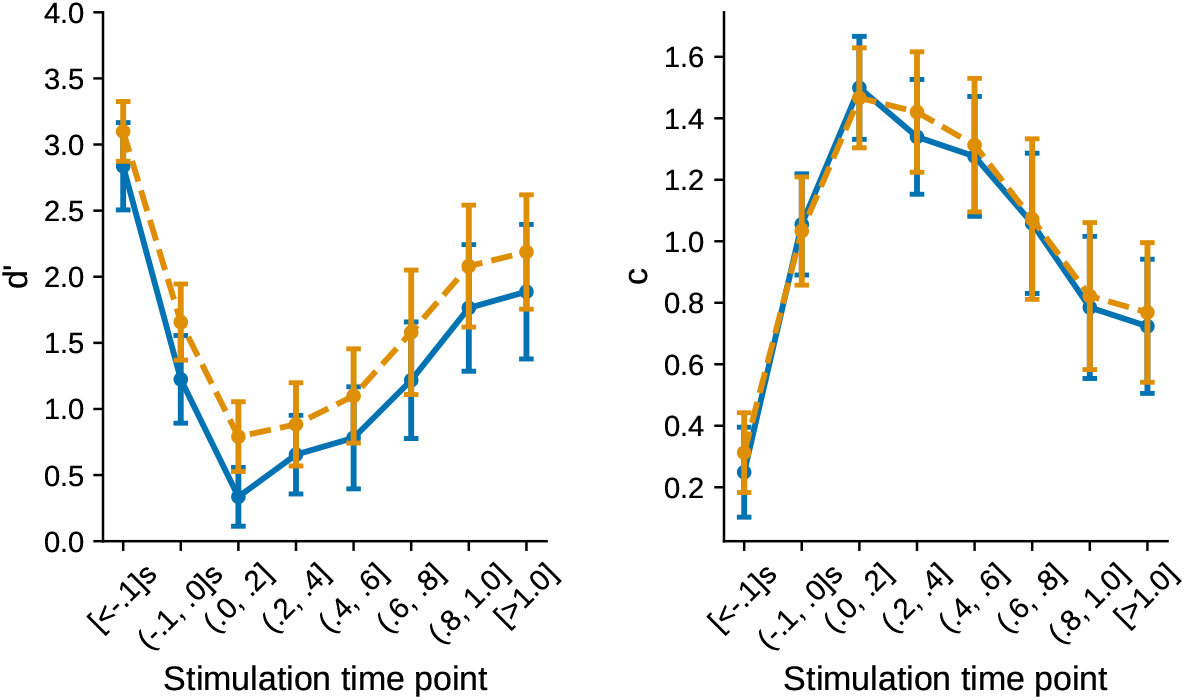
Mean and 95% confidence interval of *d*^′^ and criterion across all subjects.

### B Per subject sensitivity and criterion

**Figure S2.**
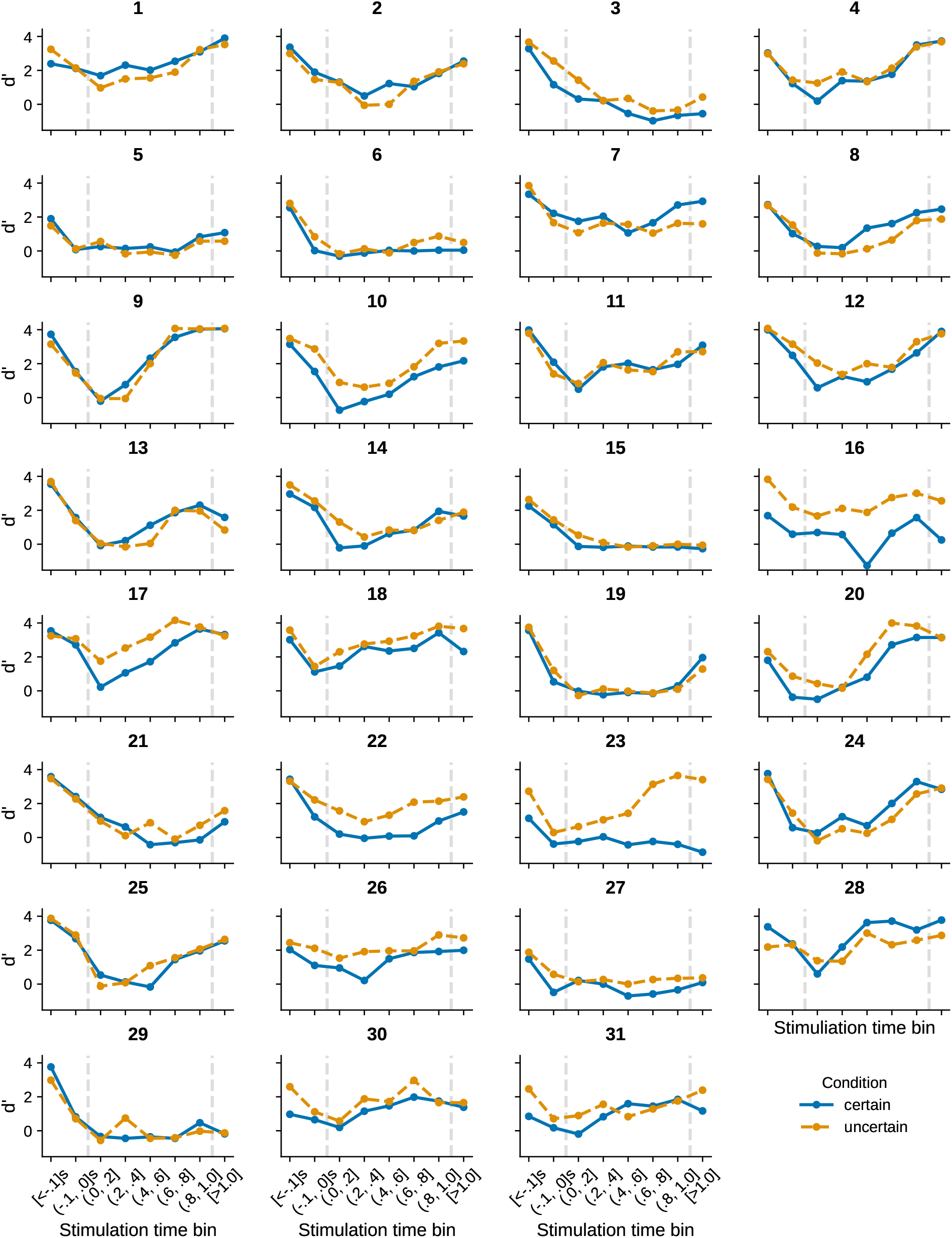
*d*^′^ per subject.

**Figure S3.**
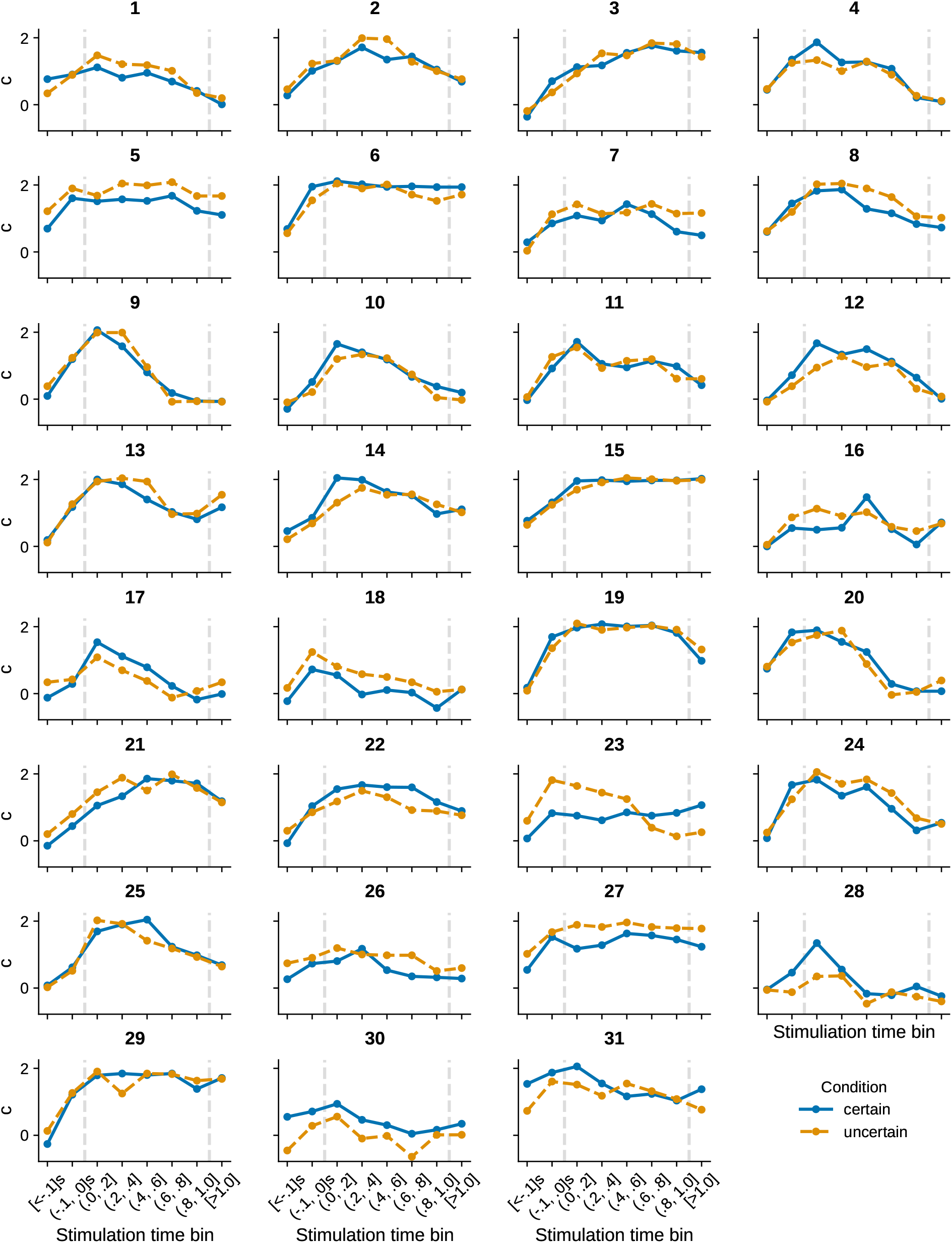
Criterion per subject

### C Per-bin paired comparison of tactile sensitivity between the *uncertain* and *certain* condition

**Table 1.**
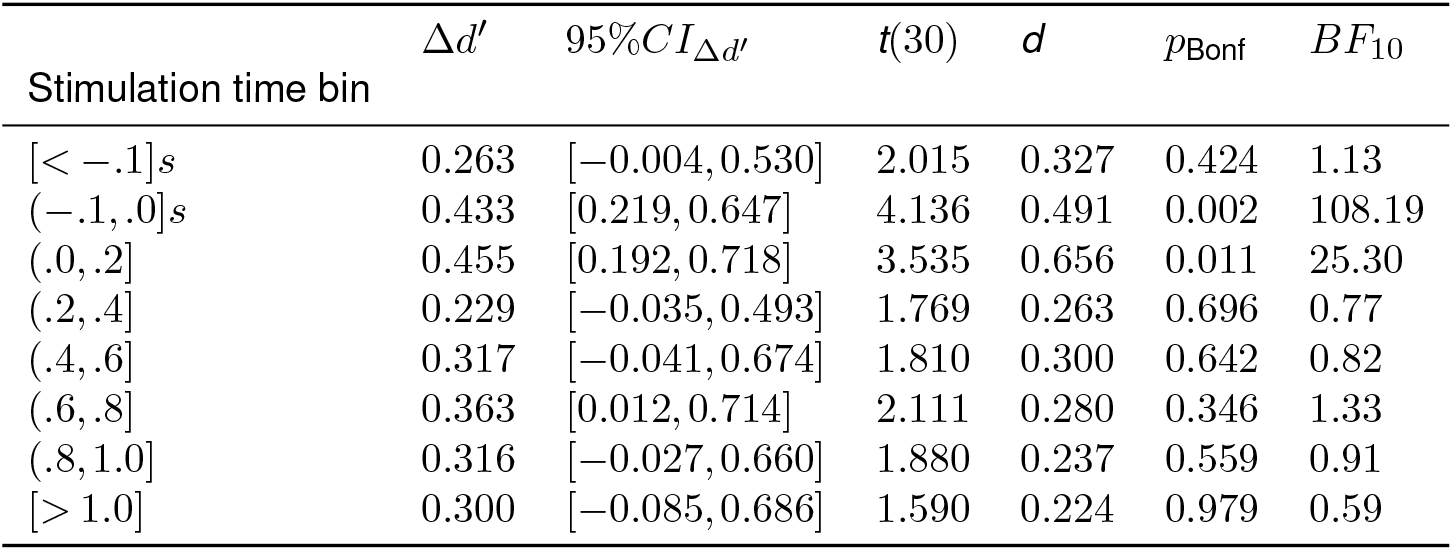
Per-bin paired comparisons of tactile sensitivity (*d*^′^) between the *certain* and *uncertain* conditions 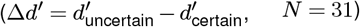, with its 95% confidence interval, Cohen’s *d*, Bonferroni-corrected *p* values, and Bayes factors *BF*_10_.

### D Kinematics normal reaching vs. initial uncertainty

**Table 2.** Kinematic summary statistics for the *certain* and *uncertain* conditions. Values are the mean across participants (*N* = 31) of each participant’s mean, with its 95% confidence interval.

| Kinematic variable | *certain* | *uncertain* |
| --- | --- | --- |
| Mean reaction time [s] | 0.287 [0.274, 0.300] | 0.284 [0.272, 0.296] |
| Mean movement time [s] | 0.671 [0.628, 0.714] | 0.691 [0.648, 0.735] |
| Mean max. velocity [m/s] | 1.538 [1.421, 1.655] | 1.490 [1.395, 1.585] |
| Mean time max. velocity [s] | 0.287 [0.263, 0.310] | 0.298 [0.274, 0.322] |
| Mean max. acceleration [m/s<sup>2</sup>] | 19.902 [16.856, 22.949] | 18.962 [16.026, 21.898] |
| Mean time max. acceleration [s] | 0.099 [0.083, 0.114] | 0.105 [0.087, 0.122] |
| Mean max. deceleration [m/s<sup>2</sup>] | 9.201 [7.535, 10.868] | 8.827 [7.276, 10.378] |
| Mean time max. deceleration [s] | 0.397 [0.369, 0.425] | 0.414 [0.381, 0.446] |

### E Optimal estimation and control

The discrete-time linear dynamical system with signal-dependent noise used to model the reaching task (Eq. 4) can be written in the following general form [1]

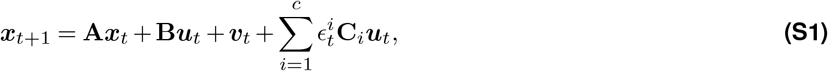

where ***v*** ∼ *N* (0, **Σ**_***v***_) and 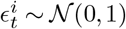 are Gaussian noise terms.

The sensory feedback (Eq. 5) can be written as

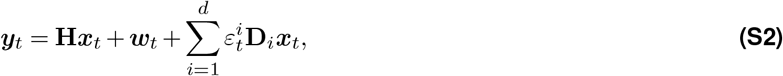

where ***w*** ∼ *N* (0, **Σ**_***y***_) and 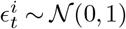 are Gaussian noise terms.

The quadratic cost function (Eq. 11) can be written as

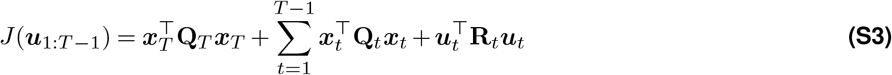

This partially observable linear system with signal-dependent noise is of the form defined by Todorov [1]. An approximately optimal sequence of filter gains **K**_1:*T* −1_ and control gains **L**_1:*T* −1_ can be found iteratively by solving for the optimal sequence of filter gains given a sequence of control gains and vice versa, and iterating that procedure until convergence (for details see Todorov [1]). The update of the state estimate then takes the following form

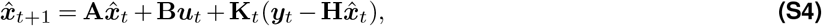

and the control input is linear in the state estimate

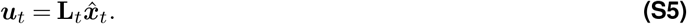

### F Model of the reaching task

We model reaching movements using a simplified model of a single-joint arm movement [1]. The hand is modeled as a point mass with position *p*(*t*), velocity 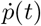, and force *f* (*t*), and an auxiliary state *g*(*t*) used to implement a muscle-like low-pass filter. We assume that the target is located at *p*^*^ = 0 and that the position *p*(*t*) is the hand’s distance to the target. In summary, the state is 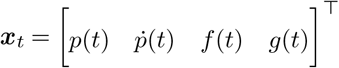 .

The discrete-time dynamics of the point mass are

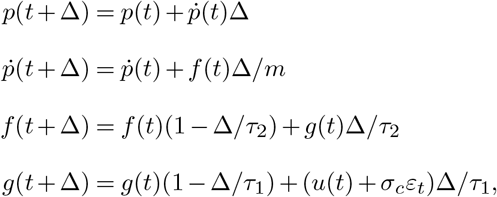

which can be represented in the form of Eq. (S1) with

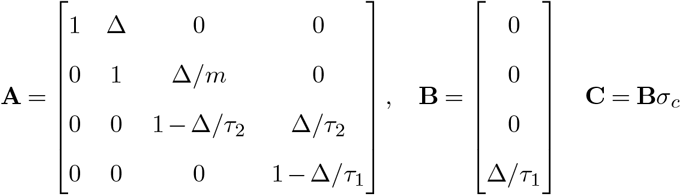

The observation matrix and sensory noise covariance are

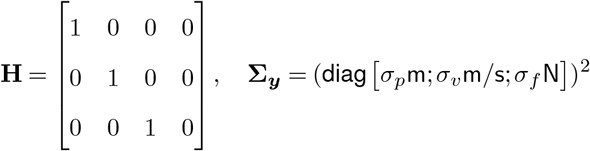

The cost function is **Q**_*t*_ = 0 for *t*_start_ *< t < T* and

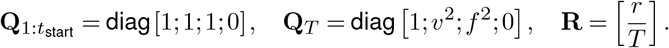

With this choice of costs we formulate the task of the participants as staying at the start position until *t*_start_, then decreasing the distance to the target at final time step *T*, with reasonable final velocity and final force, while minimizing overall energy expenditure.

In summary, the model has the following free parameters

- Mass *m* = 1.0kg
- Time constant *τ*_1_ = *τ*_2_ = 40ms
- Sensory position noise magnitude *σ*_*p*_ = 0.04
- Sensory velocity noise magnitude *σ*_*v*_ = 0.14
- Sensory force noise magnitude *σ*_*f*_ = 0.7
- Initial uncertainty in relative position *σ*_0_ = 0.0083
- Control noise magnitude *σ*_*c*_ = 0.7
- Final velocity cost *v* = 0.04
- Final force cost *f* = 0.4
- Control cost *r* = 10^−5^

We set these parameters to fit the behavioral findings of our task. But consider our analysis of robustness in Appendix H, which shows the stability of the main effect that we report about the Kalman gain over position.

### G Model simulation details

Discrete time steps were set to Δ*t* = 0.01s. The number of time steps after the GO-cue was given and before the actual movement started was set to *t*_start_ = 20. Start positions were sampled from a normal distribution 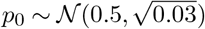 .

For the model simulations in Fig. 2f, we simulated 1000 trajectories for each participant and set the overall time to 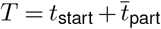, whereby 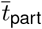 is the number of time steps closest to the individual participants’ average movement time.

For the depiction of the Kalman gains in Fig. 2d, Fig. 3c and Fig. 5c we show the Kalman gain of the model with *T* = 91 approximately corresponding to the average movement time across all participants plus the initial period.

To simulate the increased initial uncertainty in Fig. 3c and Fig. 3f, we set *σ*_0_ = 0.0093. All other model settings were identical to the previous simulation in Fig. 2f.

To account for the increased sensory uncertainty in Fig. 5c, we set *σ*_*p*_ = 0.045. All other model settings were identical to the standard parameters.

### H Influence of the model parameters on the Kalman gain over relative position

In order to investigate how the free parameters of our model influence the time course of the Kalman gain of the hand’s relative position, we varied one of the parameters while keeping all other parameters constant on the values described in Appendix F. The result of this is shown in Figure S4. We find that the cost parameters *v, f* and *r* have no or only little influence on the time course of the feedback gain. The largest influence comes from the noise parameters and the initial uncertainty in relative position. If the scale parameter of the signal-dependent motor noise *c* becomes small, sensory feedback becomes superfluous during the movement, as the forward model can precisely predict the future trajectories. When sensory velocity noise *σ*_*v*_ or force *σ*_*f*_ becomes relatively low in comparison to the sensory position noise *σ*_*p*_, the system does not need to rely on somatosensory feedback about the hand’s relative position anymore, as this can be derived from the velocity, or force information. The initial uncertainty in the belief about the hand’s relative position *σ*_0_ has a large influence on how the trajectory behaves at the start. Consider that this value will unlikely be equal to 0, as a perfectly certain belief at the beginning of a movement is unlikely due to sensory noise. Additionally, the initial uncertainty in the belief needs to be considered together with the sensory position noise *σ*_*p*_, as sensory feedback over time reduces this uncertainty. Therefore, the two values together have the biggest influence on the absolute gain about the hand’s relative position. This is why we considered experimental manipulations of those two parameters (increasing initial uncertainty in relative position and increasing uncertainty about sensory feedback). Additionally, we conducted a robustness analysis of the overall form of the time course of the Kalman gain over relative position. We computed the Kalman gains of our model for 1000 random parameter combinations of our model, where at every run we sampled each parameter uniformly in a range of 0.1 − 10 times the parameter values described in Appendix F. Figure S5 shows the average normalized Kalman gain and the standard deviation across all parameter combinations. We find that the qualitative time course we report is robust to these parameter variations on average. We note that the effect varies for different parameter combinations, but it also needs to be considered that we observe variation across different participants in our experiment.

**Figure S4.**
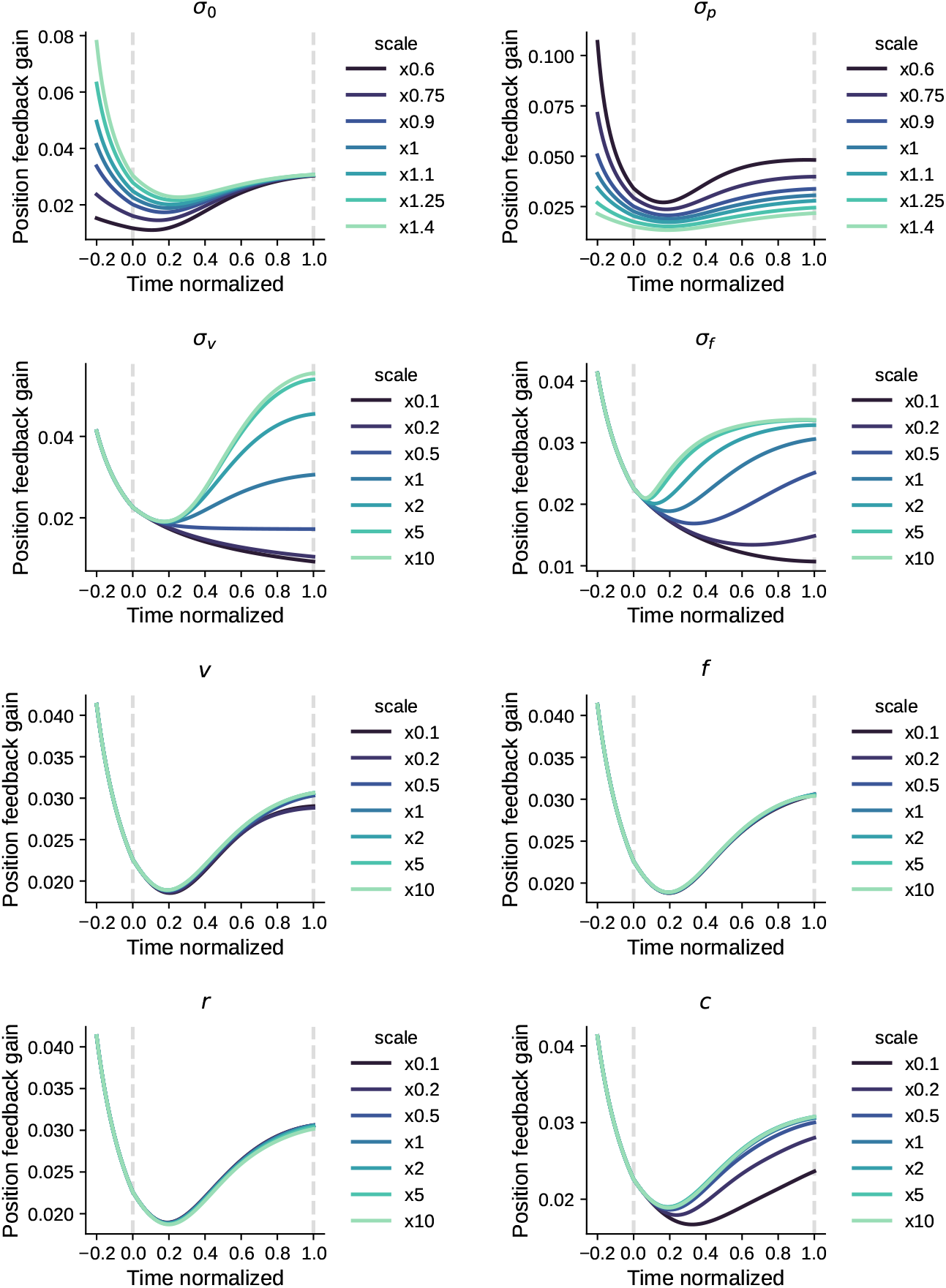
Influence of the free model parameters on the Kalman gain about relative hand position. All parameters except for the investigated parameter were kept constant. The parameter under investigation was scaled as denoted in the legend.

**Figure S5.**
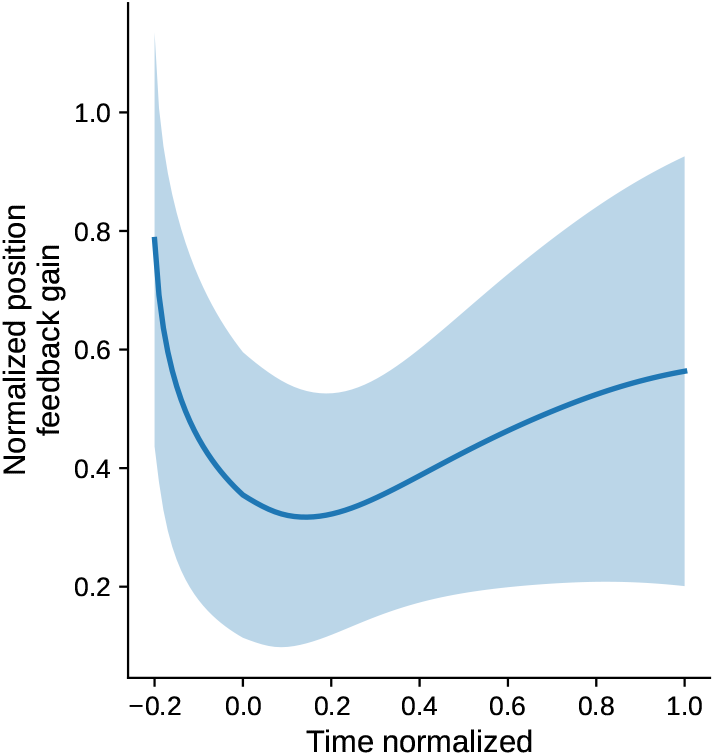
Average and standard deviation of normalized Kalman gain relative position for 1000 random parameter combinations.

**Table 3.** Model comparison on the *certain* condition. *k*: number of free parameters; *R*^2^ and MSE: goodness of fit; AIC and BIC: information criteria, with ΔAIC and ΔBIC the difference to the best model. Models are ordered by AIC and the best model is in bold.

| Model | $k$ | $R^2$ | MSE | AIC | ΔAIC | BIC | ΔBIC |
| --- | --- | --- | --- | --- | --- | --- | --- |
| **OFC K-gain** | 68 | 0.759 | 0.417 | 622.8 | 0.0 | 861.7 | 0.0 |
| Conv acc (Asym) | 62 | 0.702 | 0.515 | 663.4 | 40.6 | 881.2 | 19.5 |
| Conv acc (Sym) | 62 | 0.685 | 0.544 | 677.0 | 54.2 | 894.9 | 33.2 |
| Conv v+a (Asym) | 93 | 0.735 | 0.457 | 695.8 | 73.0 | 1022.5 | 160.8 |
| Acc-only | 62 | 0.640 | 0.621 | 709.6 | 86.8 | 927.5 | 65.8 |
| Conv v+a (Sym) | 93 | 0.719 | 0.486 | 710.9 | 88.1 | 1037.6 | 175.9 |
| Conv vel (Asym) | 62 | 0.614 | 0.666 | 727.0 | 104.2 | 944.8 | 83.1 |
| Conv vel (Sym) | 62 | 0.594 | 0.700 | 739.5 | 116.7 | 957.3 | 95.6 |
| Linear (v+a) | 93 | 0.680 | 0.552 | 742.5 | 119.7 | 1069.2 | 207.5 |
| Vel-only | 62 | 0.575 | 0.734 | 751.1 | 128.3 | 968.9 | 107.2 |

### I Model comparison detailed results

**Figure S6.**
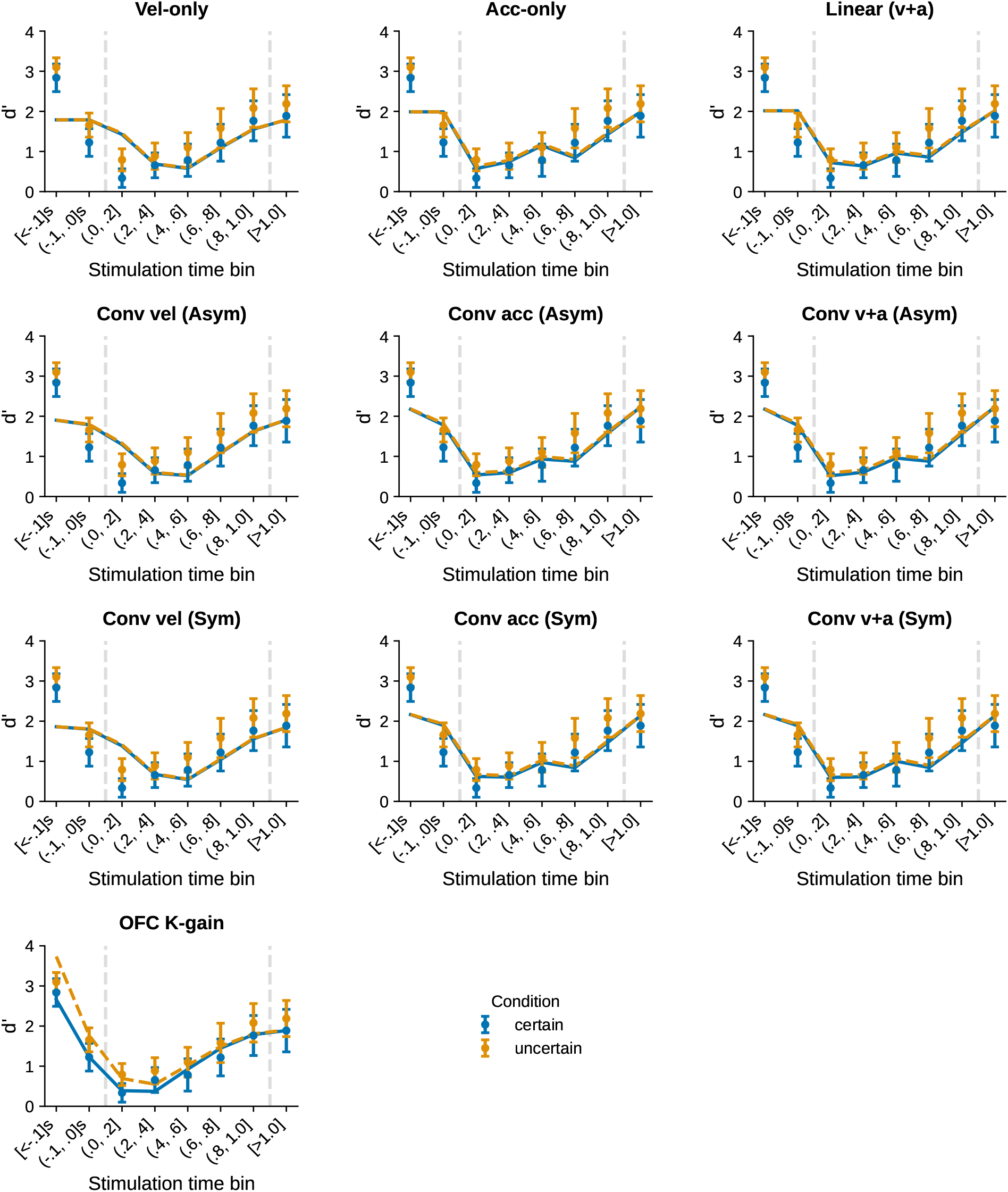
Generalization to the *uncertain* condition of the regression link on all the candidate signals fit on the *certain* condition. None of the masking models predicts a difference between the conditions. Data shows mean and 95% confidence interval.

